# Bridging DNA contacts allow Dps from *E. coli* to condense DNA

**DOI:** 10.1101/2024.01.22.576774

**Authors:** Sneha Shahu, Natalia Vtyurina, Moumita Das, Anne S. Meyer, Mahipal Ganji, Elio A. Abbondanzieri

## Abstract

The DNA-binding protein from starved cells (Dps) plays a crucial role in maintaining bacterial cell viability during periods of stress. Dps is a nucleoid-associated protein that interacts with DNA to create biomolecular condensates in live bacteria. Purified Dps protein can also rapidly form large complexes when combined with DNA *in vitro*. However, the mechanism that allows these complexes to nucleate on DNA remains unclear. Here, we examine how DNA topology influences the formation of Dps-DNA complexes. We find that DNA supercoils offer the most preferred template for the nucleation of condensed Dps structures. More generally, bridging contacts between different regions of DNA can facilitate the nucleation of condensed Dps structures. In contrast, Dps shows little affinity for stretched linear DNA before it is relaxed. Once DNA is condensed, Dps forms a stable complex that can form inter-strand contacts with nearby DNA, even without free Dps present in solution. Taken together, our results establish the important role played by bridging contacts between DNA strands in nucleating and stabilizing Dps complexes.

**Graphical Abstract:** Graphical Abstract.
Working model of nucleation and formation of Dps-DNA complex.
Regions of supercoiled or stochastically bent DNA act as nucleation points for the formation of Dps-DNA complexes by allowing Dps to form bridging contacts. Dps does not readily bind to straight stretches of DNA in isolation. Once Dps-DNA complexes are formed they can form bridging contacts to bind additional DNA.

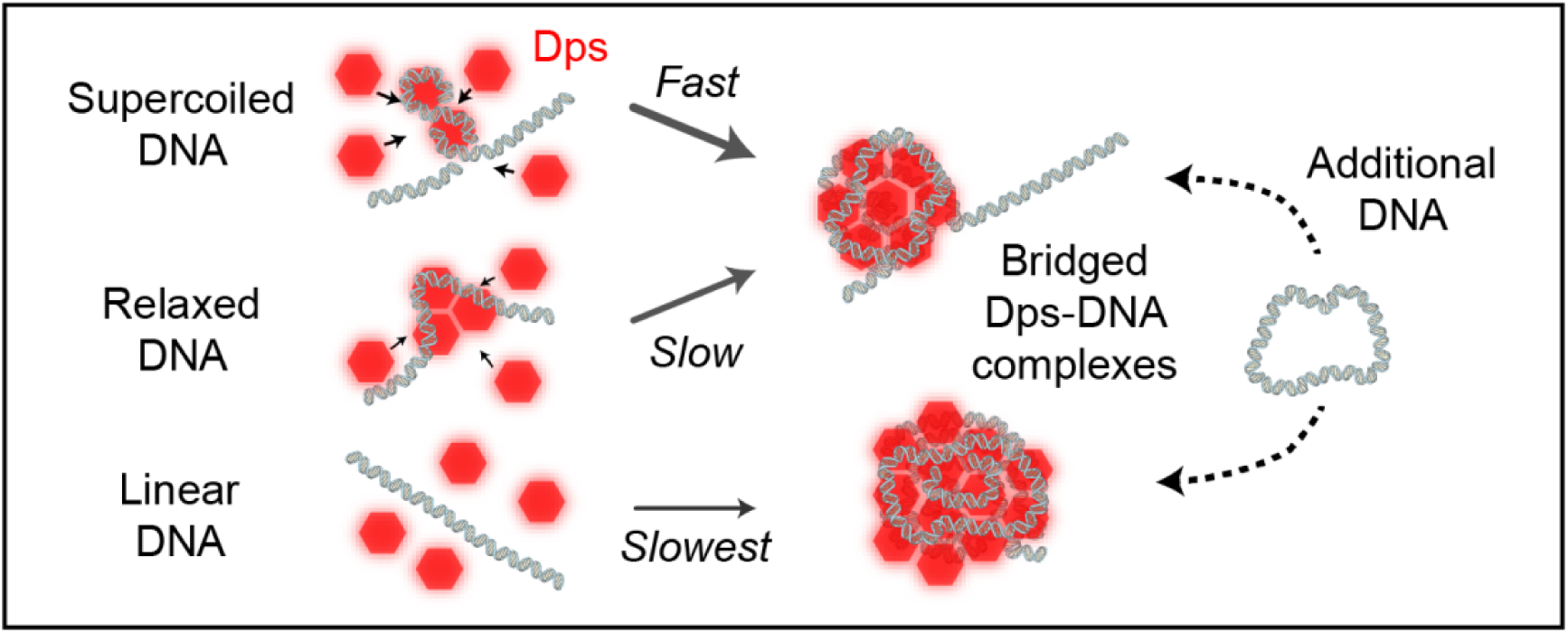

## Introduction

Nucleoid-associated proteins (NAPs) play a central role in compacting, organizing, and protecting the genomes of bacteria(1). However, the mechanisms by which NAPs interact with DNA show enormous diversity. Some NAPs, such as HU(2), IHF(3), and Fis(4), bind a single region of DNA and create a sharp bend to induce compaction. In contrast, NAPs such as H-NS(5) and DNA-binding proteins such as ParB(6) can form bridging contacts between separate regions of DNA to induce compaction. In most cases, proteins that form bridging contacts also display alternate binding modes in which they oligomerize on a single DNA strand. For example, under favorable conditions H-NS can form stiffening filaments on linear DNA instead of forming bridging contacts(5). Similarly, ParB can bind at a specific site and form linear filaments before crosslinking DNA strands with bridging contacts(6).

In stationary phase, the most abundant NAP in *Escherichia coli* is Dps (DNA-binding protein from starved cells), increasing from around 1 monomer per kilobasepair (kbp) in exponential phase to 40 monomers per kbp in late stationary phase(7). Dps both compacts chromosomal DNA and protects it from environmental stresses(8,9), including nutrient depletion, elevated temperatures, high salt concentrations, reactive oxygen species, and antibiotics(10). In *E. coli*, Dps monomers assemble into roughly spherical dodecamers with a 9 nm diameter in solution(11). The disordered N-terminal regions of the 12 Dps subunits within a dodecamer are evenly spread out around the periphery of the dodecamer and are required for tight binding to DNA(10). When these dodecamers are exposed to DNA, they bind in a sequence-independent manner(12) and rapidly condense the DNA. DNA and Dps condensates display a range of morphological features *in vivo*, suggesting that these condensates can be organized in multiple ways(13). The most well-studied morphology of Dps-DNA condensates features dodecamers organized in a regular, tightly-packed array termed a biocrystal(14). Current models of the biocrystal structure propose that Dps dodecamers make bridging contacts between parallel DNA strands, which are relatively straight(15,16), supporting a bridging mechanism for DNA compaction. However, it has also been claimed that Dps compacts DNA by bending DNA strands in a manner similar to HU, IHF, Fis(17).

We recently showed that Dps binds and compacts DNA through a cooperative mechanism described by an Ising-derived model(18). Our Ising model implies that multiple points of contact increase the avidity of Dps-DNA interactions at equilibrium. However, equilibrium models will not address how such bound complexes might nucleate and expand on a DNA strand. This nucleation of Dps condensates could occur through several possible mechanisms. Dps could induce bends in DNA which stabilize additional binding of dodecamers, similar to Fis. Alternately, Dps might form filaments on isolated strands of DNA that then form bridging contacts to other strands, similar to H-NS and ParB. Alternately, Dps may wait until two strands of DNA are in close proximity and nucleate at these sites. These points of near contact between two sections of DNA could arise from DNA packing or from supercoiling. To test these possibilities, we designed an Intercalation-induced Supercoiled DNA (ISD) assay(19) to create different DNA topologies and examined how the binding of Dps to DNA is influenced by these topologies. Using this ISD assay, we tested the Dps binding to supercoiled, bent, and linear DNA. We find that Dps preferentially binds wherever multiple DNA sections are in close proximity, such as plectonemes formed by supercoiling or locations where two DNA strands were brought together under flow. We find no evidence that Dps can stably bind on linear regions of DNA, implying that it cannot form filaments like H-NS or ParB. Finally, we see that Dps condensates can form bridging contacts to additional DNA strands in solution, even in the absence of free Dps. Our data support a model where Dps dodecamers stably bind when they are forming bridging contacts between different DNA sections.

## Materials and Methods

### Preparation of DNA and proteins

DNA molecules were prepared by digestion of plasmid pSupercos lambda 1,2 (provided by S. Hage, Delft University of Technology, Delft, The Netherlands) with XhoI (New England Biolabs) as described previously(18). Each end of the digested plasmid was ligated to a 500 bp DNA strand containing multiple biotins, resulting in a 20.6 kb DNA molecule with multiple biotins at both ends. The full-length molecules were purified from non-ligated molecules by gel electrophoresis separation on 0.8% Agarose. However, because the lengths of the ligated and non-ligated molecules were similar, this purification was not 100% efficient. With this preparation we therefore obtained DNA molecules with biotins on both the ends, only at one of the ends, or no biotins at all. Preparation of Cy5-labeled Dps was performed as previously described(18). Briefly, Dps containing a single engineered cysteine residue (T79C) was expressed and purified from *E. coli* cells, then labeled with a Cy5-maleimide dye. We typically obtained approximately 10% Cy5-labeled Dps subunits, resulting in an average of one labeled subunit in a functional dodecamer. Cy5-labeled Dps was previously shown to bind and condense DNA similarly to wild type Dps.(18)

### Single-molecule fluorescence assay to visualize different DNA topologies

Microfluidic flow cells of volume ∼10 µL were assembled by sandwiching double-sided tape between a quartz slide and a cover slip. The inner surfaces of the flow cell were passivated with a 20% (w/v) solution of PEG and biotin-PEG at a 100:1 ratio dissolved in a freshly made 100 mM bicarbonate buffer, preventing non-specific adhesion of proteins and DNA to the surface. We then applied 20 µL of a 0.1 mg/mL streptavidin solution into an empty flow cell and incubated for 1 minute. Unbound streptavidin was removed by flowing 200 µL of buffer T50 (50 mM Tris⋅HCl, pH 8.0, and 50 mM NaCl) through the flow cell. Biotin-labeled DNA of around 10 pM was then applied onto the PEG/biotin-PEG surface at a constant flow rate of 20 µL/min to immobilize the DNA via biotin-streptavidin-biotin interactions. Since the biotin-labeled DNA contained biotin handles on either one side or two sides, we obtained both singly-tethered and doubly-tethered DNA molecules. Excess unbound DNA and any DNA that did not contain biotins on either end were removed during the buffer wash. Labeled or wild-type Dps was diluted to a final concentration of 25-200 nM in imaging buffer (40 mM Tris-HCl pH 7.3, 50 mM KCl, 20-50 nM Sytox Orange (SxO), 5% PEG 8K, 2 mM Trolox, 40 µg/ml glucose oxidase, 17 µg/ml catalase, and 5% glucose) and introduced to the flow cell to induce compaction. We washed flow cells with 500 µL of imaging buffer, after which undigested SxO-labeled pSupercos lambda 1,2 plasmid DNA was introduced by adding 100 µL of 10 pM plasmid DNA suspended in imaging buffer.

We used a previously described(20) home-built fluorescence microscope for direct observation of different topological DNA molecules and their interactions with Cy5-labeled Dps. With a 60x water immersion objective (Olympus UPLSAPO, 1.2 NA), we imaged the sample through a narrow slit (Thorlabs) to remove light originating from outside a narrow band within the image plane. We then split the path into the two emission colors. Having the slit and separate paths for each color allowed us to display and record both colors on an electron-multiplying charge-coupled device (EMCCD) camera (Andor Ixon 897) simultaneously. Further details are provided in the supplementary information.

For imaging DNA compaction by wild type Dps at different concentrations, a Nikon Ti2 Eclipse microscope equipped with a motorized H-TIRF, perfect focus system and a Teledyne Photometrics PRIME BSI sCMOS camera were used. Illumination for the visualization of DNA was done with a 561 nm wavelength laser using an L6cc laser combiner (Oxxius Inc., France). Imaging was done with an oil immersion objective lens (Nikon Instruments Apo SR TIRF 100×, numerical aperture 1.49, oil) under HiLo TIRF illumination.

### Bootstrap analysis

Because the condensation times were not normally distributed, we chose to use a bootstrapping approach to determine if the null hypothesis could explain the differences between DNA strands grouped by different topologies(21). To compare two sets of data, we pooled the sets into a single data set (simulating the assumed null distribution) and then randomly drew 10^7^ simulated pairs of data sets from this pooled data set. Absolute values of the differences in the mean of the simulated paired sets were compared to the observed difference, allowing us to robustly estimate the two-tailed *p*-value.

### MCMC simulations

Markov Chain Monte Carlo (MCMC) simulations were performed using Igor Pro software. The DNA strand of 20.6 kb was divided into 343 Dps binding sites of equal affinity, with each binding site 60 bp in length. At each step, a binding site was chosen at random and toggled between bound or unbound. The change in free energy was then calculated based on three modeled scenarios: non-cooperative binding, cooperative nearest-neighbor interactions, and an extended cooperative binding model that allows up to six nearest neighbors to interact on each side of the binding site. If the change in free energy were negative, the change was accepted. If the free energy increased, a Gibbs term was used to calculate the probability that the change was accepted as:

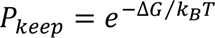

where *k*_*B*_ is the Boltzmann constant and *T* is temperature. A pseudo-random number generator was then used to create a number between 0 and 1, and the change was accepted if this result were lower than *P*_*keep*_. For further details see supplementary methods. Additional details of the MCMC simulations are described in the supplementary information.

## Results

### A double-tether assay produces three DNA topologies

Previously we have described a technique to introduce supercoiling into doubly-tethered DNA molecules using intercalating dyes, an assay we named Intercalation-induced Supercoiled DNA (ISD)(19). Here we take advantage of the ISD assay to visualize various DNA conformations. We introduced a linear 20.6 kb DNA strand that had been stochastically labeled with multiple biotins at either one or both the ends into a streptavidin-coated flow cell at a constant flow rate of 20 µL/min (Fig. 1A-top). After one end of the DNA attached to the surface, the DNA strand was stretched in the direction of applied flow. For strands with biotin labels at both ends, the other end of the DNA also attached to the surface, creating a doubly-tethered DNA strand (Fig. 1A). The doubly-tethered DNA molecules were stretched on average to 52% of the contour length. We visualized the DNA molecules using epi-fluorescence microscopy by staining with Sytox Orange (SxO), which unwinds the DNA slightly as it intercalates between base pairs.

**Figure 1:**
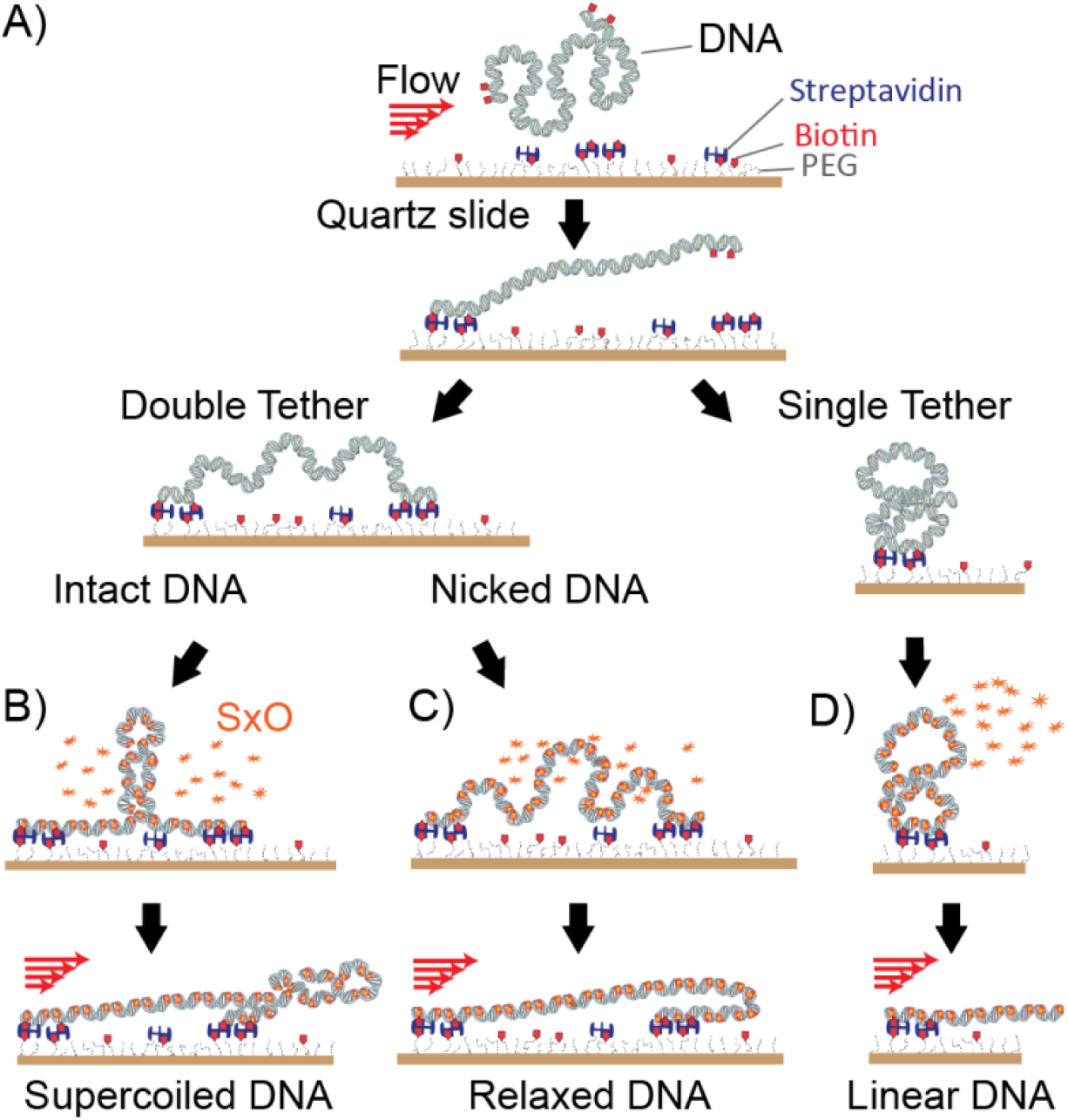
Generation of three different DNA conformations. A) (top) DNA with biotins at one or both ends are introduced into a streptavidin-coated flow cell at a constant flow rate. (bottom) During the flow, DNA binds to the surface at one end and becomes stretched in the direction of the flow. If biotin is present at the distal end, then the DNA forms an extended double tether, otherwise it remains singly tethered. B) Upon addition of SxO, doubly-tethered intact DNA becomes supercoiled. C) Nicked DNA remains torsionally relaxed. D) Singly-tethered DNA is torsionally relaxed. Upon application of flow, the molecules from B)-D) assume readily identifiable shapes.

Once SxO had intercalated into the DNA, we observed plectonemes induced by supercoiling on some fraction of doubly-tethered DNA molecules(19) (Fig. 1B). These molecules adopted a Y-shape upon application of flow (Fig. S1A). Another fraction of doubly-tethered DNA molecules was torsionally relaxed (e.g. through a nick in the backbone) and did not exhibit plectonemes (Fig. 1C). These molecules became J-shaped under flow (Fig. S1B). Before flow was applied, supercoiled DNA exhibited dynamic bright fluorescent spots corresponding to the plectonemes while relaxed DNA showed uniform intensity and a greater flexibility, allowing them to be distinguished(19,22). Finally, in the same field of view we also obtained molecules that were tethered at a single end (Fig. 1D) and displayed a restricted diffusive motion around the tethered position. These molecules became stretched in the direction of applied flow, adopting an I-shape (Fig. S1C). Since all DNA molecules studied were tethered within the same flow cell, they experienced identical buffer conditions over time. Therefore, any statistically significant differences in their behavior could be directly related to the topology of DNA.

### Dps preferentially nucleates condensation at parallel DNA strands

We used single-molecule total internal reflection fluorescence (TIRF) microscopy to directly visualize the binding of Dps to different conformations of DNA. Cy5-labeled Dps was introduced to the flow cell while alternating laser excitation at 532 nm and 642 nm was used to monitor the shape of the DNA and the accumulation of bound Dps. We flowed in Cy5-labeled Dps at a concentration of 200 nM and a flow rate of approximately 600 µL/minute. Dps was observed to arrive at the imaged DNA approximately 5 seconds after the flow had begun (Vid. S1). As described above, DNA molecules adopted one of three conformations during the flow: supercoiled and doubly tethered (Y-shape), relaxed and doubly tethered (J-shape), or relaxed and singly tethered (I-shaped) (Fig. 2 A-C). Among the singly-tethered molecules, we also identified a handful of instances where the flow brought the DNA strand into close proximity to a downstream DNA strand, allowing for inter-strand contacts to form (Fig. 2D).

**Figure 2.**
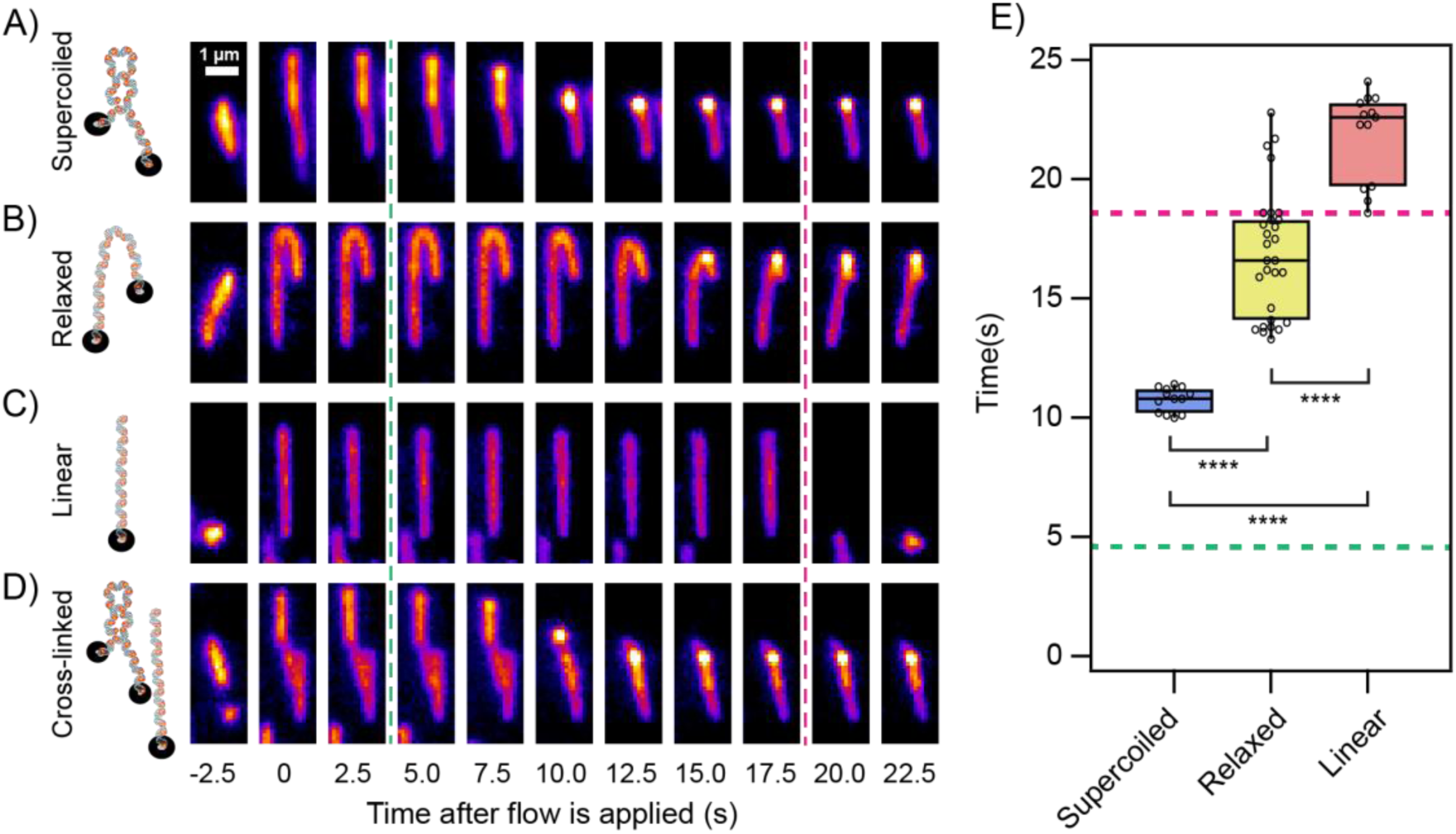
Interaction of Dps with different DNA topologies. **(A-D)** Diagram showing DNA of different topologies and snap-shots of corresponding fluorescence images before and during application of flow. **(A)** Supercoiled DNA becomes Y-shaped under flow. **(B)** Diagram of relaxed DNA and corresponding fluorescence images showing that the DNA becomes J-shaped under flow. **(C)** Diagram of singly-tethered DNA and its corresponding fluorescence images showing DNA becoming stretched due to flow. **(D)** Diagram of a singly-tethered DNA molecule and a doubly-tethered DNA exhibiting plectonemes. Corresponding fluorescence images show the two strands are not contacting each other before arrival of Dps. The two molecules were cross-linked by binding of Dps. **(E)** Boxplot showing the distribution of timepoints when Dps fully compacted the DNA in each conformation. Note that a considerable number of relaxed DNA molecules and all the linear molecules were fully compacted only after flow ceased. All images were acquired at a 100 msec frame rate. Flow began immediately after frame 1. Free Dps arrives approximately 6 seconds later (green dashed line). Flow was halted after 20 seconds (red dashed line).

We measured the amount of time it took for the compaction of each molecule to finish. We have included snapshots of the compaction of representative molecules from each category (Fig. 2 A-D), and we have plotted kymographs showing compactions as a function of time (Fig. S2). Compaction typically began at a single nucleation site that increased in fluorescence intensity as more DNA and Dps was added to the Dps-DNA complex. Compaction was observed to cease either when the entire strand was compacted (in the case of singly-tethered DNA molecules) or when the remaining DNA was stretched tautly (in the case of doubly tethered molecules). Supercoiled DNA (Y-shaped) was compacted the most rapidly, with compaction nucleating on the plectoneme immediately after Dps arrived and proceeding at a near constant rate for ∼6 seconds until compaction was complete (Fig. 2A, Fig. S2A, Vid. S2). Relaxed DNA that was doubly tethered (J-shaped) took longer to nucleate compaction, but once compaction began it proceeded at a similar rate (Fig. 2B, Fig. S2B, Vid. S3). Nucleation typically occurred near the point of maximum curvature. Not all J-shaped molecules completed compaction before the flow was stopped. Once the flow ceased, compaction was detected by the formation of a bright spot along the DNA strand and a loss of Brownian fluctuation in the remaining DNA. Singly-tethered DNA (I-shaped) was unable to nucleate compaction in isolation during flow, but compaction could be detected after flow ceased by a decrease in the mobility of the DNA(18). All of these molecules eventually compacted when the flow was halted after 20 seconds. Compaction times were determined for each category of molecule and were compared (Fig. 2E). These results demonstrate that the topology of DNA has a significant effect on the kinetics of compaction by Dps.

To further confirm that DNA topology affects the kinetics of compaction, we repeated these experiments using various concentrations of wild type Dps and a milder flow rate (Fig. 3). Flow was applied for 30 seconds at a rate of 100 µL per minute and Dps was introduced at concentrations ranging from 25 nM up to 200 nM. Once the flow was stopped, the molecules were observed for an additional 180 seconds to detect slower compaction events. At 25 nM Dps, we observed compaction of supercoiled DNA but failed to measure compaction of relaxed or singly-tethered DNA, even once the flow was halted. These topologies are either unable to support compaction at all or compact over timescales much longer than the experiment. At higher Dps concentrations, we observed an identical trend in the mean compaction times as observed using labeled Dps, with supercoiled DNA compacting first, relaxed DNA compacting second, and singly tethered DNA compacting last. We also performed an experiment using a lower SxO concentration (20 nM) with a Dps concentration of 50 nM, producing a lower supercoiled density in the non-nicked DNA strands because of fewer sites of intercalation. We once again observed an identical trend in compaction times between the three categories (Fig. S3).

**Figure 3.**
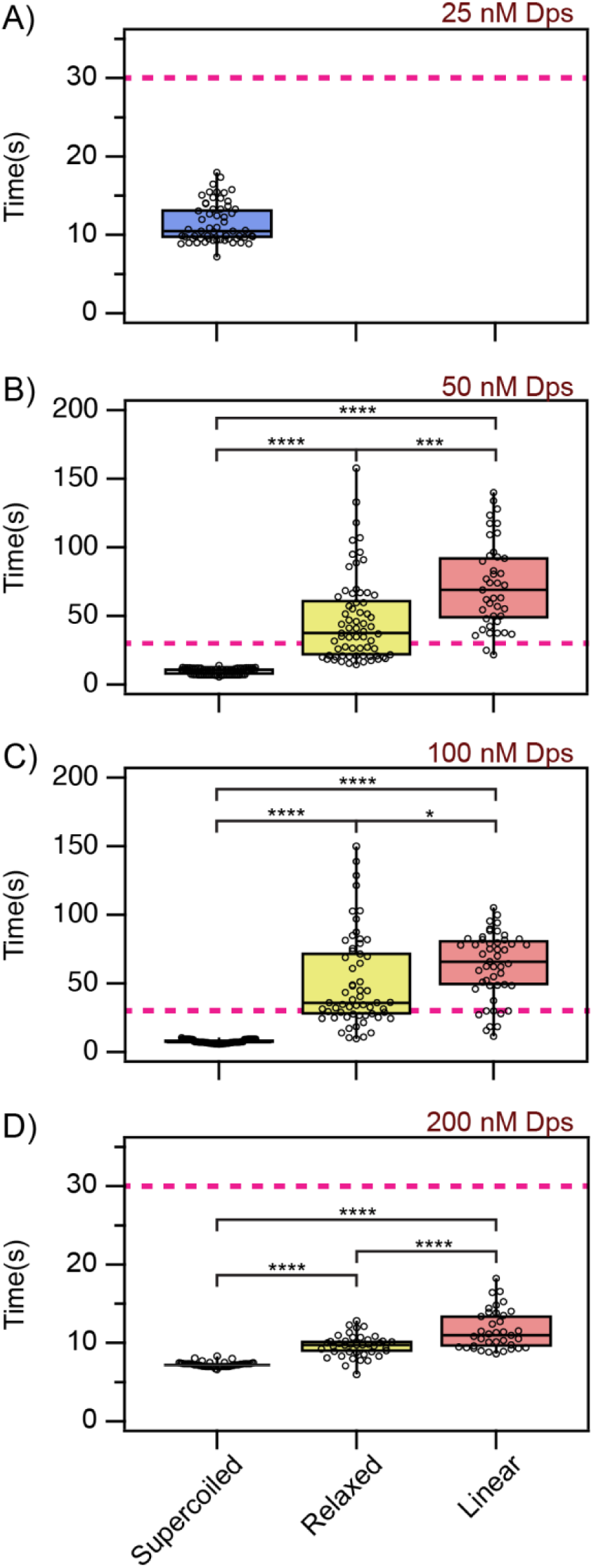
Boxplots showing DNA compaction times for a range of unlabeled Dps concentrations. **(A)** Flow at 100 µL/minute was halted after 30 seconds (red dashed line). Boxplots showing the distribution of timepoints when 25 nM unlabeled Dps fully compacted the DNA in each conformation. No compaction events were observed for relaxed or single-tethered (linear) molecules. **(B)** Same plat as in **(A)** but at 50 nM unlabeled Dps. **(C)** Same plat as in **(A)** but at 100 nM unlabeled Dps. **(D)** Same plat as in **(A)** but at 200 nM unlabeled Dps.

For comparison, we also measured the effects of introducing a buffer containing 20% PEG 8k and 3 mM MgCl_2_ to different DNA topologies. These concentrations of PEG and Mg^2+^ have been shown to condense DNA in single molecule measurements using magnetic tweezers, similar to Dps(23). In our assay, we did not observe any condensation by PEG under flow, presumably due to the shearing forces applied on the DNA. After the flow ceased, some DNA strands condensed while other strands remained locked in an extended conformation (Fig. S4) with no observable preference for supercoiled or relaxed topologies.

### Dps can mediate cross-linking between DNA strands

Because of the high density of immobilized DNA in our assay, at several locations we could observe interaction of two DNA molecules in the presence of Dps. Interestingly, we found that Dps was capable of mediating the cross-linking of two initially independent DNA molecules (Fig. 2D). In the example shown, the upper molecule was doubly tethered and supercoiled while the lower DNA strand was singly tethered. During the flow, the singly-tethered DNA molecule was stretched (0-10 s) over the doubly-tethered molecule. Dps nucleated and compacted the doubly-tethered molecule, and this compact structure cross-linked to the singly-tethered molecule (12.5 s), decreasing its Brownian motion. Cross-linking was apparent after the flow was stopped (22.5 s) and the singly-tethered DNA remained stretched rather than relaxing to its original position. A kymograph demonstrating the cross-linking of two linear DNA strands is also shown (Fig. S2D).

### Bound Dps is distributed unevenly on doubly-tethered DNA

Our previous study using single-molecule techniques revealed that reducing the tension applied to DNA via magnetic tweezers causes Dps to abruptly compact the molecule through a cooperative Ising mechanism(18). Singly-tethered DNA molecules showed this behavior in our flow assay as well, with a rapid collapse of the molecule beginning immediately upon the cessation of flow. However, for a doubly-tethered molecule, compaction will result in increasing tension on the DNA arising from the connections to the coverslip. As a portion of the DNA is condensed, the remaining DNA must increase its extension relative to its contour length because the total distance between the tether points remains unchanged. Applying the well-established worm-like chain model for the force-extension relationship of DNA(24,25), this extension necessarily requires the tension in the molecule to increase. We investigated the effect of this increased tension on the final distribution of Dps on these molecules by measuring the fluorescence intensity of both the DNA and Dps reporters once the applied flow was stopped.

Prior to the introduction of Dps, doubly-tethered molecules were highly dynamic and showed either diffusive fluctuations of plectonemes for supercoiled DNA or lateral fluctuations in position for relaxed DNA. After the flow was stopped and Dps had bound the DNA, we no longer observed fluctuations across the DNA length in doubly-tethered molecules (Fig. S1), indicating that these DNA molecules were held under tension due to compaction. This finding is consistent with our previous measurements that a tension of greater than ∼2 pN will stall compaction by Dps under similar conditions(18). We imaged the DNA-Dps complexes using prism-based TIRF microscopy with sensitivity sufficient to detect single fluorophores(20). DNA-Dps complexes can be readily identified as overlapping bright fluorescence spots from SxO (DNA, green) and Cy5 (Dps, magenta) (Fig. 4A). Strikingly, Dps was only detected in a single mass at one end of the DNA and not observed along the length of the stretched DNA. This was true both for molecules that condensed under constant flow and for molecules that condensed after the flow had ceased, indicating that the observed singular distribution was not due to the applied flow.

**Figure 4.**
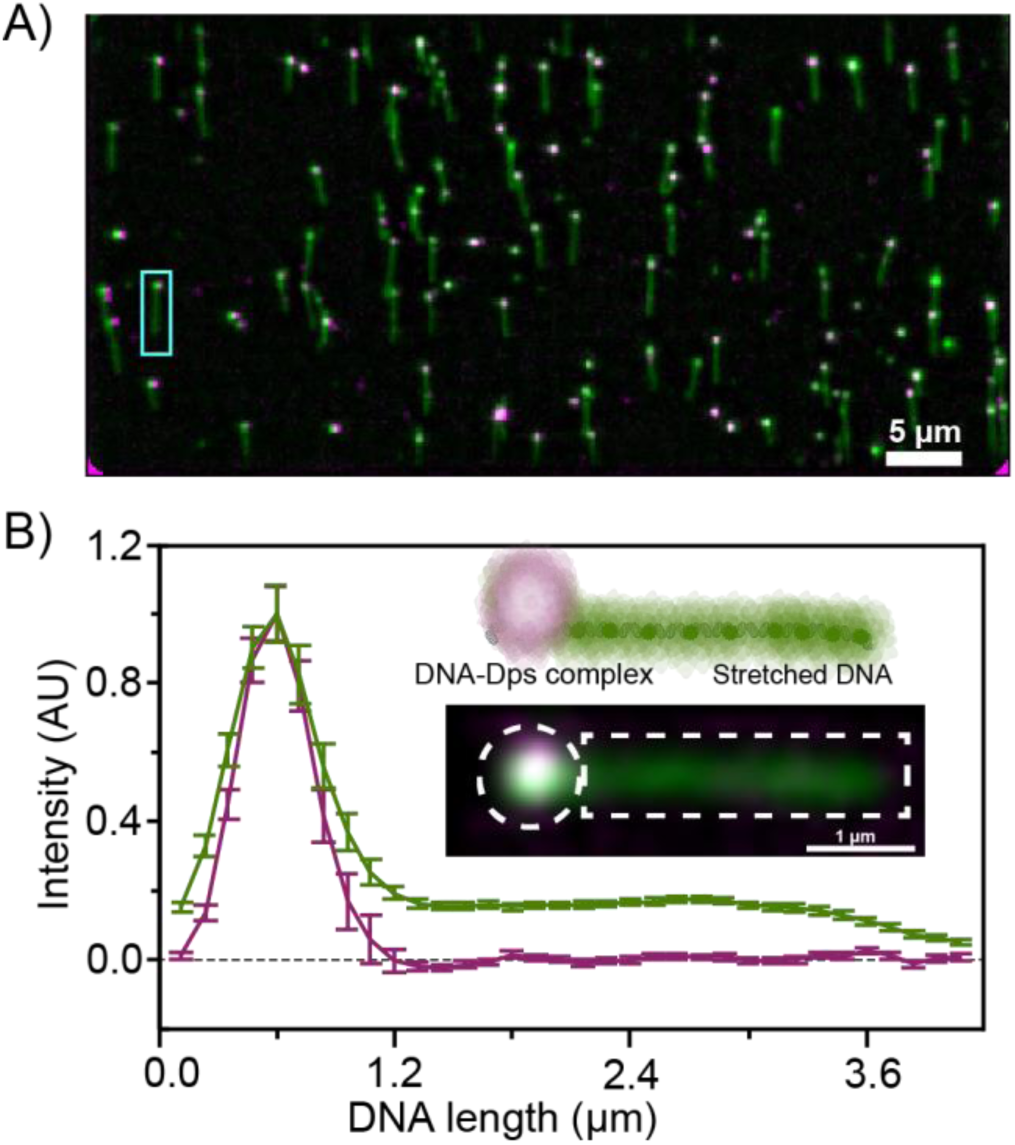
Distribution of DNA and Dps molecules along doubly-tethered DNA. **(A)** Overlaid fluorescence signals obtained from DNA labeled with SYTOX orange (green) and Cy5-labeled Dps (magenta) after the formation of Dps-DNA complexes and the cessation of flow. A representative example of a stretched DNA molecule with a Dps-DNA complex is highlighted (cyan rectangle). **(B)** Plot showing the intensity distribution of Cy5-Dps (magenta) along DNA labeled with SYTOX orange (green) obtained by averaging the intensity of Cy5 and SxO from 90 different structures. Inset represents the example molecule from **(A),** illustrating orientation of pixels used in the plot.

We then performed quantitative analysis of the distribution of Dps along doubly-tethered DNA molecules. An example molecule for analysis is indicated with a rectangle in Fig. 4A. The fluorescence intensities corresponding to DNA (SxO) and Dps (Cy5) from 90 individual molecules were averaged along the length of the DNA-Dps complexes (Fig. 4B). We found no detectable Dps signal outside of the single diffraction-limited condensates at one end of the complexes. The DNA was also concentrated in the condensate, which accounted for around 50% of the SxO signal in each molecule. Strikingly, we never saw instances where multiple condensates separated by stretched DNA formed on a single molecule. These results demonstrate that Dps has a strong bias to bind condensed DNA, taking advantage of the increased avidity and cooperative interactions associated with these structures.

### Markov Chain Monte Carlo (MCMC) simulations establish that a modified Ising model is consistent with the measured distribution of Dps along DNA molecules

The observation that Dps formed one condensate per DNA molecule that did not extend beyond a diffraction-limited spot, rather than binding at several locations along the DNA, suggests that Dps binds in a highly cooperative manner. To further investigate this relationship, we conducted MCMC simulations to test three possible scenarios for the degree of cooperativity between Dps dodecamers (see Methods).

In the first scenario, we assumed that Dps dodecamers bind independently to the DNA. In the second scenario, we assumed that dodecamers are more weakly attracted to the DNA but are stabilized by nearest-neighbor interactions if the DNA on either side is occupied and compacted by Dps. In the third scenario, inspired by the cooperative Ising model we developed previously**(18)**, we assumed that bound dodecamers can be stabilized by interactions from up to six nearest neighbors on each side of the binding site. In each case, we also fixed the total extension of the DNA and used an analytic approximation of the extensible worm-like chain model**(26)** to calculate the energy needed to stretch the unbound DNA to this extension. We chose values for the binding energy and the energy of nearest-neighbor interactions in each scenario to ensure that approximately 50% of the DNA was bound at equilibrium (Table S1). We then ran simulations for a sufficiently large number of steps to ensure that the system had arrived at a stable minimum energy and analyzed the binding pattern of Dps.

In the non-cooperative scenario, the amount of bound Dps approached equilibrium rapidly (Fig. S5) and was nearly uniformly distributed across the DNA (Fig. 5a). We simulated the distribution of the fluorescent density on such a molecule (see Methods) and found that Dps density would be unevenly but continuously distributed across the DNA molecule (Fig. 5b). In the nearest-neighbor scenario, the system took slightly longer to reach equilibrium (Fig. S5) and Dps was less evenly distributed, but multiple Dps clusters always formed (Fig. 5c). The simulated fluorescence density distribution showed multiple distinct peaks of Dps (Fig. 5d). In the Ising scenario that allowed for long-range interactions, the simulation took longer to reach equilibrium (Fig. S5) but always settled into a single strong Dps peak (Fig. 5e). The simulated fluorescence densities also displayed this single strong peak of Dps (Fig. 5f) and bore a strong resemblance to the measured density (Fig. 4b). A similar pattern was observed when the binding energies were reduced by 50% in all three scenarios (Fig. S6). These simulations therefore support a model where Dps dodecamers interact with many binding sites along the DNA. Because the stoichiometry of Dps bound to DNA is estimated to be 60 base pairs per dodecamer**(27)**, interactions across multiple binding sites imply that individual dodecamers are either crosslinking distal DNA regions or crosslinking to other dodecamers bound to distal DNA regions.

**Figure 5.**
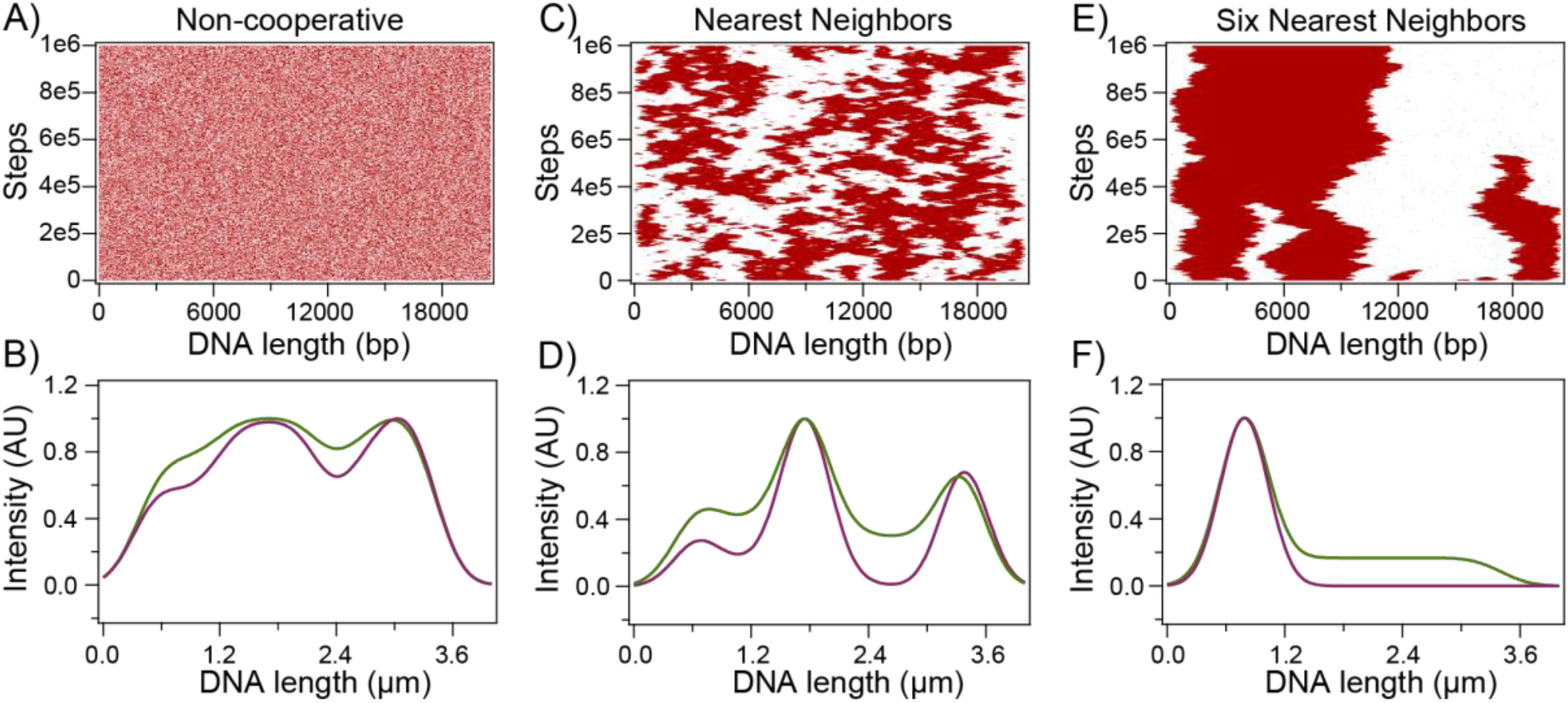
Markov Chain Monte Carlo (MCMC) simulations of Dps;DNA complex formation. **(A)** An MCMC simulation was performed where each Dps 12-mer was attracted to the DNA by a chemical potential of 6 k_B_T for binding DNA but with no cooperative interactions between neighboring 12-mers. Red dots represent DNA bound to Dps, and white patches represent unbound DNA. **(B)** Simulated fluorescence plot of a representative distribution of Dps (red) and DNA (green) seen in the simulation from (A). **(C)** A simulation as in (A) but with 0 k_B_T chemical potential for binding DNA and a 6 k_B_T interaction between any 12-mers bound in adjacent sites. **(D)** Simulated fluorescence plot of a representative distribution of Dps (red) and DNA (green) seen in the simulation from (B). **(E)** A simulation as in (A) but with 0 k_B_T chemical potential for binding DNA and a 1 k_B_T interaction between up to six adjacent 12-mers. **(F)** Simulated fluorescence plot of a representative distribution of Dps (red) and DNA (green) seen in the simulation from **(E)**.

### Stable Dps-DNA complexes can be formed at a variety of stoichiometries

We next aimed to determine whether the stability of Dps-DNA complexes would be affected by changing the ratio of DNA to Dps. We therefore used the preformed Dps-DNA condensates formed at Dps concentrations ranging from 50 nM to 200 nM as a base to bind additional DNA. To ensure that little to no free Dps remained in the solution, we thoroughly washed the flow cell with 10-50 times its volume of buffer. As expected, the preformed Dps-DNA complexes remained stable and compacted even after free Dps was depleted, consistent with our previous findings that Dps dissociates slowly once a stable complex is formed(18).

Next, we introduced undigested 10 pM SxO-labeled plasmid DNA (20.6 kB) to the flow cells while monitoring the fluorescence of DNA from the preformed Dps-DNA complexes. Additional DNA was injected and flow cells were monitored over a period of 6 minutes. The plasmid DNA flowing in the buffer formed mobile spots larger than the diffraction limit, indicating that the DNA remained flexible with no Dps bound (Fig. 6A). Occasionally, an unbound plasmid DNA strand would encounter a preformed Dps-DNA complex and rapidly bind to it, as shown in panels 9 and 10 of Fig. 6A. Once such an additional plasmid strand was captured, the plasmid DNA condensed to a diffraction-limited spot. The fluorescence from the SxO channel at that spot increased sharply (Fig. 6B), resulting in the ratio of Dps to DNA in the fused complex decreasing by 50% or more. After the step-like increase, the fluorescence intensity remained stable, indicating that a new equilibrium had been reached. We also tested the ability of linear DNA to bind preformed Dps-DNA condensates and observed identical behavior as plasmid DNA.

**Figure 6.**
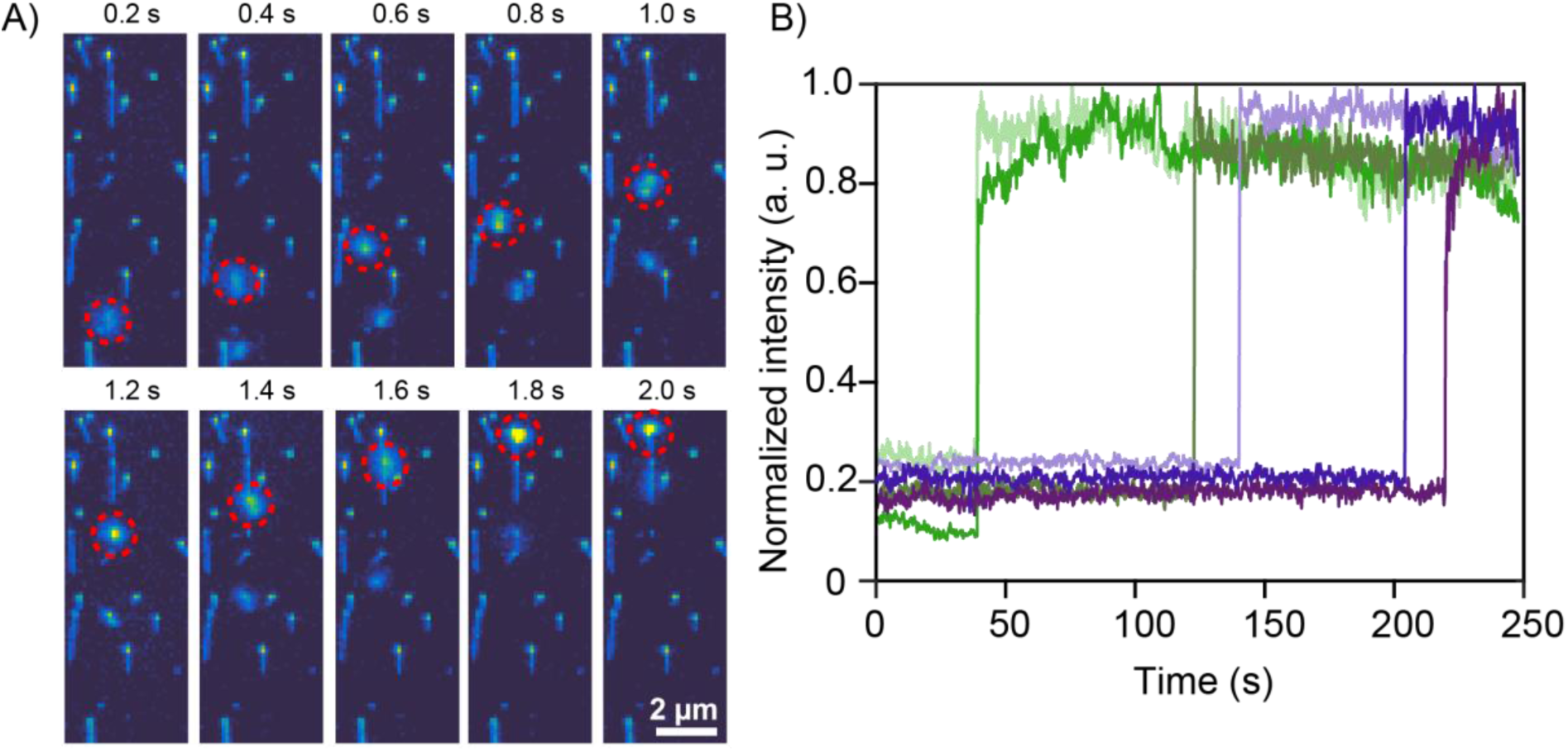
Binding of DNA plasmid to preformed Dps-DNA assemblies. **(A)** DNA plasmid (shown in red circle) freely moving towards a preformed Dps-DNA assembly (frame 1-8) and binding to it (frame 9 and 10). Each frame is a snapshot taken at 200 msec intervals. **(B)** Normalized time traces of six events where a single-step sharp increase in DNA fluorescence intensity of the DNA-Dps complex indicated binding of an extra DNA plasmid.

To measure the efficiency of the capture process 100 preformed condensates were selected at random from each field of view and the fraction of complexes that bound additional DNA was calculated (Fig. S7). We observed that complexes formed at a low initial Dps concentration (50 nM) were less likely to capture additional DNA strands (∼5% efficiency), while complexes formed at higher initial Dps concentrations (100-200 nM) were better able to capture additional DNA strands (∼20% efficiency). These findings demonstrate that Dps can condense DNA even if many potential binding sites on the DNA remain unbound, indicating that Dps and DNA can form stable complexes over a range of stoichiometries. The existence of complexes containing multiple DNA strands also provides additional direct evidence that Dps can crosslink different strands of DNA together.

## Discussion

In this report, we investigated how different DNA conformations and supercoiling states affect Dps binding, nucleation, and subsequent DNA compaction by the nucleoid associated protein Dps. Utilizing the ISD assay, we demonstrated that Dps will bind most readily to plectonemic DNA, less readily to bent or flexible DNA, and shows no detectable affinity for linear stretched DNA. When bound, Dps prefers to form a single large complex on extended DNA strands rather than forming multiple smaller complexes. Finally, we observed that Dps complexes can stably bind DNA at multiple stoichiometries. All these observations support a model in which Dps dodecamers bind in a highly cooperative manner to DNA, relying on multiple contacts to increase their stability on the DNA.

### The kinetics of compaction by Dps are strongly influenced by DNA topology

In our previous work, we showed that Dps can condense isolated DNA strands rapidly, even when under tension(18). Based on the high degree of cooperativity demonstrated by this collapse, we inferred that Dps dodecamers derive their stability from a high avidity associated with multiple DNA contacts. We also observed that the first step of collapse was rate limiting, with the DNA strand remaining fully extended for a stochastic length of time extending up to hundreds of seconds followed by a rapid binding of hundreds of dodecamers in under one second, causing it to condense.

Our current work allows us to examine in more detail the mechanistic origin of this first binding step. Our data are consistent with a model where Dps dodecamers prefer to form bridging contacts between either two distant regions of the same DNA strand that have folded back on each other or points of contact between separate DNA molecules, producing regions where two DNA strands are separated by distances similar to the diameter of a dodecamer (∼9 nm). Under these conditions, the rate of nucleation is considerably higher than for isolated strands of DNA, as evidenced by the higher overall rate of compaction on supercoiled DNA compared to relaxed or single-tethered DNA (Figs. 2, 3, and S3). In relaxed and single-tethered DNA, the probability of stochastic contacts between different regions of the DNA will increase once flow has ceased and the molecules explore a greater range of conformations through diffusion, creating favorable conditions for nucleation. Once nucleation occurs, additional condensation has a low dependence on topology, as Dps acting on both supercoiled and relaxed DNA requires around 5 seconds to compact several kilobases of DNA under flow (Fig. S2), presumably because the nucleated complex will always be in close proximity to any bare DNA strands directly adjacent to it.

This preference for bridging contacts is consistent with the structure of the Dps dodecamer. Dps binds DNA primarily through lysine-rich N-terminal regions(10). These regions are largely unresolved in crystal structures(11), but their structure has been predicted using AlphaFold AI software(28,29). An AlphaFold structural prediction for the *E. coli* Dps subunit (UniProt ID: D3QNM8) indicates that 12-13 N-terminal amino acids may be intrinsically disordered. To help visualize how Dps could potentially bind to a DNA strand, we superimposed 12 copies of the N-terminal structure predicted by AlphaFold onto the corresponding locations in the crystal structure of a Dps dodecamer (Fig. 7a). Given the size of the dodecamer (∼9 nm diameter) relative to the maximum length of this disordered region (∼4 nm), only a minority of these binding regions would be able to interact with a straight strand of DNA. Bringing a second strand of DNA close to the dodecamer would therefore double the avidity of binding, exponentially enhancing stability. In support of this model, a recent EM study observed potential nucleation structures of Dps-DNA biocrystals that featured Dps dodecamers “zipping together” parallel DNA strands(30).

**Figure 7.**
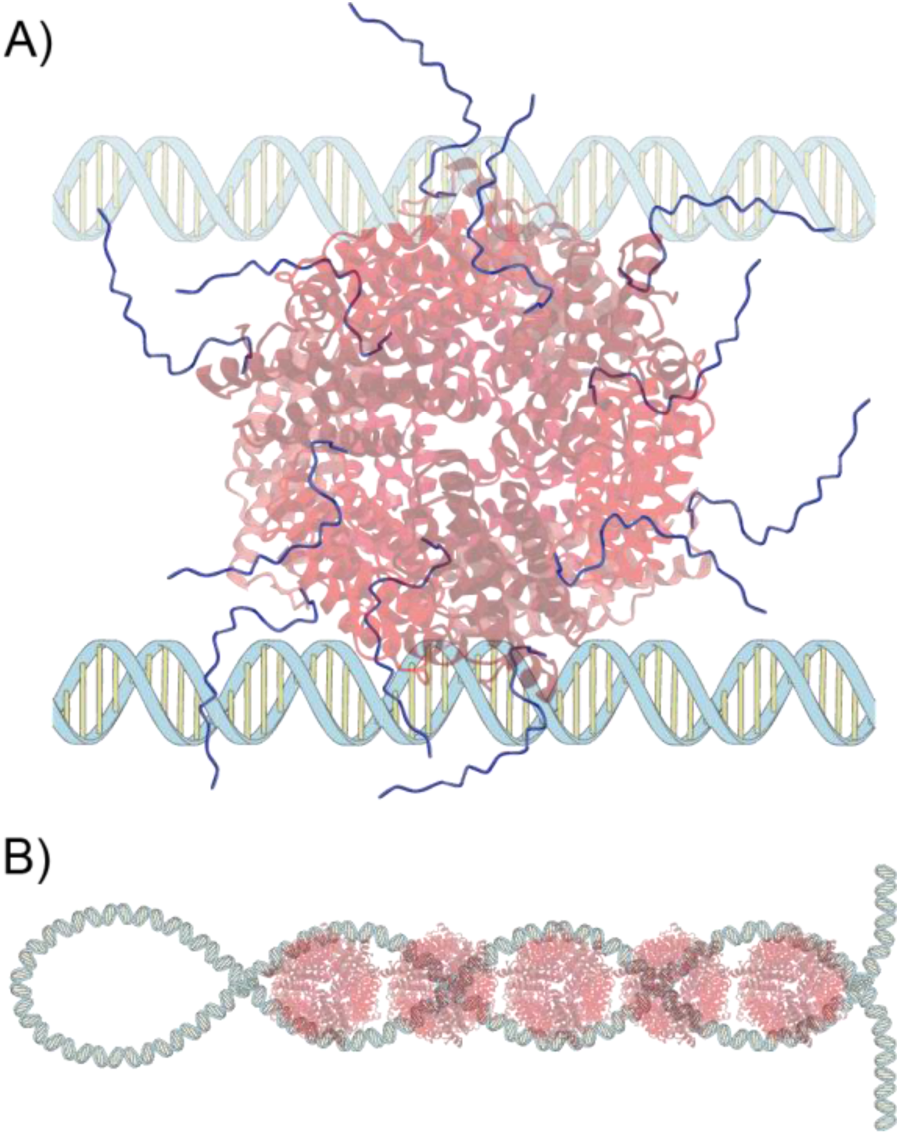
Parallel DNA strands increase Dps avidity. **(A)** A Dps dodecamer core (red) is surrounded by 12 flexible N-terminal regions (blue) that bind DNA. The large diameter of the dodecamer (∼9 nm) prevents all of the N-terminal regions from accessing a single strand of DNA simultaneously (bottom). A second strand of DNA allows many more of the N-terminal regions to bind DNA. **(B)** Models of DNA plectonemes suggest that the two DNA strands forming a helical coil are spaced by roughly 8-10 nm, allowing a single dodecamer to interact with both strands.

Our data demonstrate that plectonemes in particular offer an efficient template for Dps to bind, presumably because this topology places distant regions of a DNA strand next to each other (Fig. 7b). Indeed, theoretical models of the plectoneme structure(31) suggest that the two strands are spaced by 7-11 nm when the plectoneme is under a physiological range of tensions (1-3 pN). This spacing closely matches the spacing of DNA observed in Dps biocrystals (8-9.8 nm)(15). The size of Dps dodecamers is therefore well-suited for binding inside the helical stems of DNA plectonemes.

Under conditions of stress, the distribution of supercoiling in the bacterial nucleoid undergoes large rearrangements(32). Based on our results, this reorganization could influence the binding of Dps to the bacterial nucleoid. In support of this hypothesis, it was reported that Fis inhibits the ability of Dps to condense DNA in log phase in part by downregulating the expression of Topoisomerase I and DNA gyrase, limiting the formation of new supercoils. In contrast, overexpression of Topoisomerase I and DNA gyrase was observed to enhance condensation by Dps(33). Further research is needed to elucidate how supercoiling influences Dps-induced condensation of DNA *in vivo*.

### Dps requires multiple contacts to stably bind DNA

While we found that two DNA strands in close proximity help nucleate the condensation of DNA by Dps, on stretched regions of DNA with no possibility for bridging contacts we saw no detectable binding of Dps dodecamers (Fig. 4B). This behavior stands in contrast to observations of other proteins known to bind and condense DNA. The mitochondrial transcription factor A (TFAM), which is responsible for condensing the mitochondrial nucleoid and is known to use cooperative binding, has been shown to coat linear DNA before it condenses it(34). Similarly the Fused-in-Sarcoma (FUS) protein, a canonical model of liquid-like protein condensates, was shown to coat DNA before condensing it(35). Finally, the pioneer transcript factor Sox2 was shown to coat DNA at forces too high to allow condensation(36). The authors also observed that Sox2 forms multiple clusters on a single DNA strand that was doubly tethered to a surface. This binding pattern contrasts with our observation that Dps always formed a single complex on doubly-tethered DNA and would be consistent with Sox2 having a lower cooperativity than Dps (Fig. 5). These comparisons suggest that Dps is one of the most cooperative proteins capable of condensing DNA.

### Dps-DNA complexes can accommodate a range of stoichiometries

Our observation that preformed Dps-DNA complexes can stably accommodate additional DNA strands without absorbing additional Dps (Fig. 6, Fig. S7) demonstrates that the stoichiometry of DNA-Dps complexes is not fixed. The change in stoichiometry could occur because some DNA binding domains were not bound to DNA in the original condensate, allowing them to bind a nearby strand. Alternately, nearly all the binding domains could be engaged in the preformed condensate, but they dynamically bind and release DNA, allowing dodecamers to rapidly rearrange to accommodate an additional DNA molecule. Dps is known to be capable of forming biocrystals with regular spacing both *in vitro* and *in vivo(14)*. However, these crystalline arrays have a fixed ratio between Dps and DNA. Our results therefore require either that Dps and DNA do not always combine into regular biocrystals or that regular biocrystals can stably exist within a larger structure that includes portions of unbound DNA. In support of the former explanation, multiple morphological forms of Dps-DNA complexes have been observed *in vivo,* with clear evidence of the biocrystal morphology only observed in cells overexpressing Dps*(13)*. These results imply that the stoichiometry of Dps and DNA can affect the morphology. Allowing preformed Dps-DNA condensates to bind additional DNA may induce Dps to convert between different morphologies by altering the stoichiometry. Further research is needed to understand how the stoichiometry and morphology of Dps-DNA condensates relate to each other.

## Acknowledgments

We are grateful to Julie Biteen for fruitful discussions. We would like acknowledge the technical support from Venkat Reddy Dadireddy and Shankar Krishna. Funding to M.D., A.S.M., and E.A.A. was provided by the National Science Foundation via MODULUS DMS-2031180 and by the National Institutes of Health via 1R01GM143182-01. Funding to M.G. and N.V. was provided by the Netherlands Organization for Scientific Research (Frontiers of Nanoscience program). M. G. gratefully acknowledges funding from DST-SERB via Startup Research Grant (SRG-2021-0001553) and DBT/Wellcome India Alliance intermediate fellowship (IA/I/21/2/505928). This research was supported in part by the National Science Foundation under Grant No. NSF PHY-1748958 and the Gordon and Betty Moore Foundation Grant No. 2919.02.

## Data Availability Statement

Custom-written Igor scripts used for MCMC simulations will be provided upon request to the Lead Contact, Elio Abbondanzieri.

## Notes

### Competing Interest Statement

The authors have declared no competing interest.

