## Supplementary Information for "Bridging DNA contacts allow Dps from *E. coli* to condense DNA"

### Supplementary Data

#### Total internal reflection fluorescence (TIRF) microscopy setup

We combined two color lasers (532 nm Cobolt Samba and 642 nm Cobolt MLD) using a dichroic mirror (FF635-Di01-Semrock) and directed the resulting beam through an acousto-optic tunable filter (AOTF) from AA Optoelectronics to allow for fast alternating laser excitation (ALEX) between the two colors. This imaging modality was necessary to track two labels (SxO-labeled DNA and Cy5-labeled Dps). We then focused the beam into a prism at a near-normal incidence angle to facilitate co-localization of both colored beams at the image plane. Total internal reflection occurred at the quartz-water interface, enabling TIRF illumination of the sample. Using a 60x water immersion objective (Olympus UPLSAPO, 1.2 NA), we imaged the sample through a narrow slit (Thorlabs) to eliminate light originating from outside a narrow band within the image plane. We then split the path into the two emission colors. The use of the slit and separate paths for each color allowed us to simultaneously display and record both colors next to each other on an electron-multiplying charge-coupled device (EMCCD) camera (Andor Ixon 897).

The microscope was controlled using a custom LabVIEW routine. The timing of ALEX and EMCCD was achieved through LabVIEW NIDAQ-mx using a PCIe-6320 card and a BNC-2120 breakout box from National Instruments. In a typical experiment, we recorded approximately 2000 frames with a 100-millisecond frame rate. The first frame was obtained under the excitation wavelength of 642 nm, followed by 10 frames of 532 nm illumination. We used Tetraspeck beads (Invitrogen) to identify the paired coordinates of the red and green spots and fit the data to a fifth-order polynomial function. Mapping was necessary to co-localize Cy5-labeled Dps spots in the red channel with the corresponding SxO-labeled DNA fluorescence in the green channel.

### Markov Chain Monte Carlo simulations

Markov Chain Monte Carlo (MCMC) simulations were performed using Igor Pro software. The algorithm used was an extension of the mean field approximation used previously(1). Using an estimate of one Dps dodecamer per 60 bp(2), the 20.6 kilobase DNA strand was divided into 343 Dps binding sites. At each step, a binding site was chosen at random and toggled between bound or unbound. The new free energy was then calculated based on one of three modeled scenarios: non-cooperative binding, cooperative nearest-neighbor interactions, and an extended cooperative binding model that allows up to six nearest neighbors to interact on each side of the binding site. Energies used to model the binding energy and nearest neighbor interactions in each scenario are given in Table S1 below. For every occupied site, the total energy was lowered by the binding energy term. Nearest neighbor energies were calculated by counting the number of contiguously bound sites up to the maximum interaction distance and multiplying by the interaction energy, then subtracting this from the total free energy. Bound sites were assumed to be completely condensed, requiring us to shorten the contour length of the remaining DNA. The work needed to stretch the remaining DNA by this amount was calculated by integrating a numerical approximation of the extensible worm like chain force-extension relationship(3).

The total free energy $G_{tot}$ of a given bound state was therefore calculated as:

$$G_{tot}(b_1, b_2 \dots b_N) = -G_{BE} \sum_{n=1}^{N} b_n - G_{NN} \sum_{m=1}^{M} \sum_{n=1}^{N-m} b_{m,n} + E_{WLC}(L_C)$$

Here $b_n$ is the bound state of the nth binding site (set to 0 or 1), $G_{BE}$ is the binding energy, $G_{NN}$ is the interaction energy, $E_{WLC}$ is the work needed to stretch the remaining unbound DNA, and $L_C$ is the contour length of the unbound DNA. The index variable $b_{m,n}$ tracks if the binding site is the start of a contiguous bound patch of length m and is calculated by a recursive formula:

$$b_{m,n} = b_{m-1,n} \cdot b_{m-1,n+1}$$

$$b_{1,n} = b_n$$

Once all the energetic terms had been summed, the new energy was compared to that of the previous step. If the change in free energy were negative, the change in the bound state was accepted. If the free energy increased, a Gibbs term was used to calculate the probability that the change was accepted:

$$P_{keep} = e^{-\Delta G / k_B T}$$

A pseudo-random number generator was then used to create a number between 0 and 1, and the change was accepted if this result were lower than $P_{keep}$. Simulations were performed for 10^6 steps, which was long enough to see the total free energy plateau to a minimum level.

| Scenario tested | **1: Non-cooperative** | **2: Nearest neighbor only** | **3: Ising model (6 nearest neighbors)** |
|---|---|---|---|
| $G_{BE}$ | 6 kT | 0 kT | 0 kT |
| $G_{NN}$ | 0 kT | 6 kT | 1 kT |
| $M$ | 0 | 1 | 6 |

**Table S1**: Energy terms used in MCMC scenarios


1. Vtyurina, N.N., Dulin, D., Docter, M.W., Meyer, A.S., Dekker, N.H. and Abbondanzieri, E.A. (2016) Hysteresis in DNA compaction by Dps is described by an Ising model. *Proc Natl Acad Sci U S A*, **113**, 4982-4987.
2. Ceci, P., Cellai, S., Falvo, E., Rivetti, C., Rossi, G.L. and Chiancone, E. (2004) DNA condensation and self-aggregation of Escherichia coli Dps are coupled phenomena related to the properties of the N-terminus. *Nucleic Acids Res*, **32**, 5935-5944.
3. Petrosyan, R. (2017) Improved approximations for some polymer extension models. *Rheologica Acta*, **56**, 21-26.

**Figure S1**

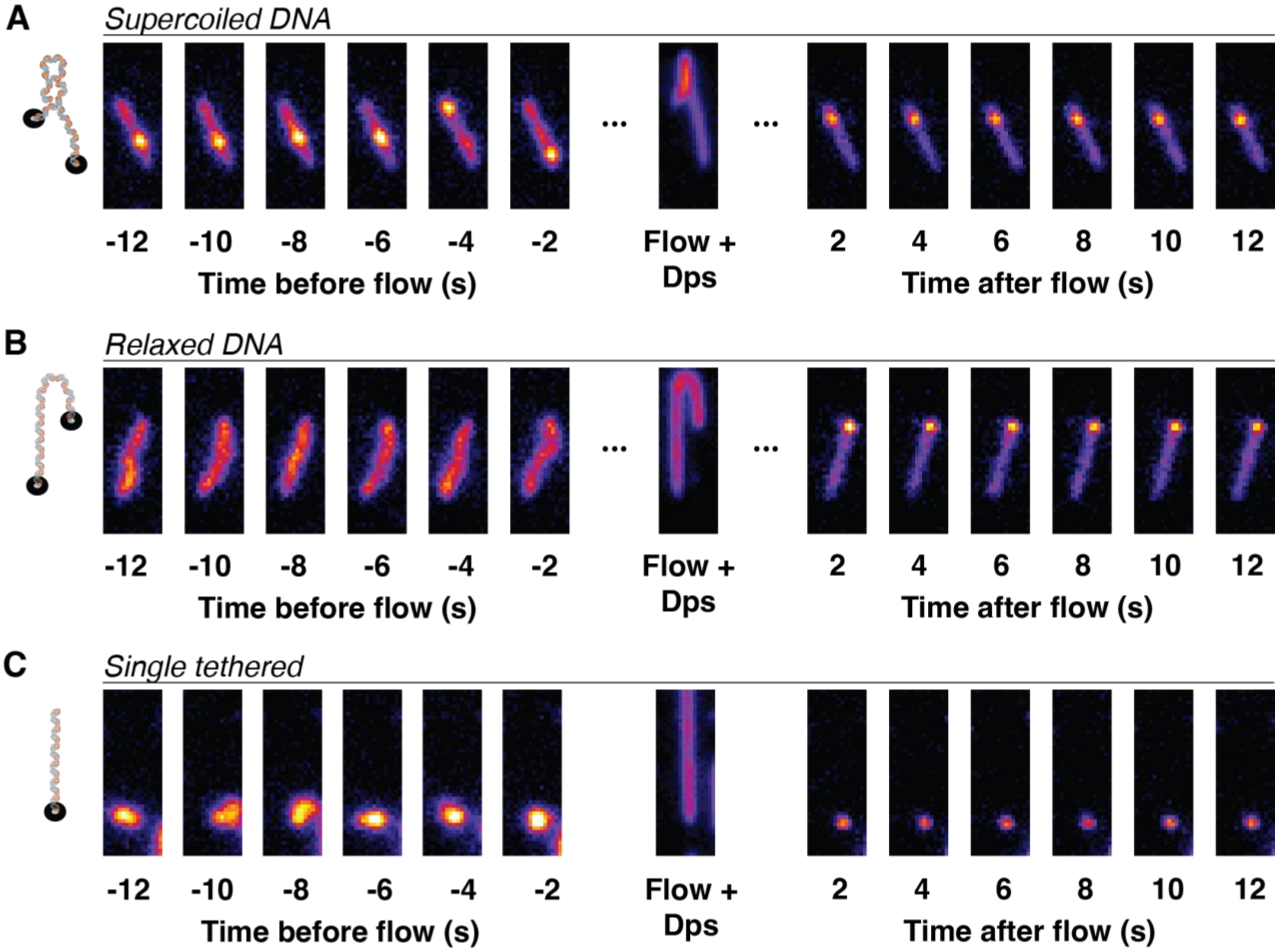


**Figure S1. Characteristic diffusive motion of different DNA topologies. (A)** A doubly-tethered DNA strand with supercoiling formed a plectoneme, which appeared as a small bright spot under zero flow. The plectoneme removed excess slack in the remaining DNA, causing the DNA to form a relatively straight line between the points of attachment. Successive frames show that the plectoneme can diffuse along the DNA with a preference for specific positions. Under flow, the plectoneme was stretched out, creating a Y-shaped molecule. After binding Dps it formed a "lollypop" structure. **(B)** A doubly-tethered DNA strand that is torsionally relaxed formed a bent and winding path between the points of attachment. Successive frames show various bent conformations. Under flow, the DNA formed a J-shaped molecule. After binding Dps it formed a "lollypop" structure. **(C)** A singly-tethered DNA strand formed a compact spot that exhibits constrained diffusive motions about a point. Under flow, the DNA was stretched out linearly creating an I-shaped molecule. After binding Dps, it formed a small spot that did not move noticeably.

**Figure S2**

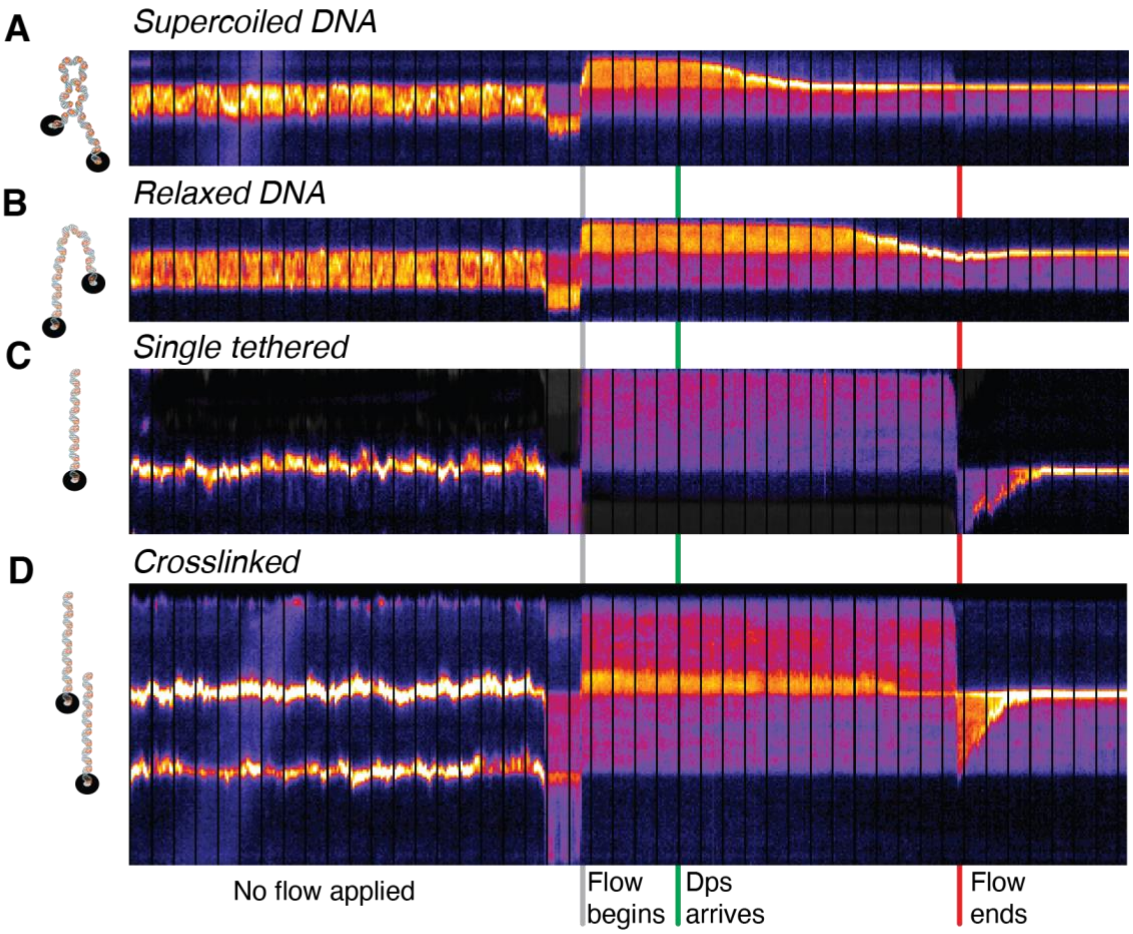


**Figure S2. Kymographs of DNA compaction. (A)** The SxO fluorescent signal shows a doubly-tethered DNA molecule with supercoiling exhibiting plectoneme dynamics before the flow is applied. After a brief period of reverse flow, the application of forward flow stretched the molecule out. Soon after Dps arrives, compaction began and proceeded for around 6 s. Once compaction was complete, the distribution of DNA was fixed. Vertical black stripes correspond to alternating excitation used to monitor Dps levels and are spaced by 1.1 s. **(B)** A relaxed DNA molecule showed a more uniform density distribution before flow is applied. Under forward flow, the molecule did not begin to compact for around 7 s. Once compaction began it proceeded for around 6 s. **(C)** A singly-tethered molecule exhibited small excursions around a fixed point of attachment before flow began. Once flow was applied, the molecule was stretched out and did not compact. After the forward flow was stopped, a brief period of weak reverse flow began during which the molecule compacted in around 4 s. **(D)** Two neighboring DNA molecules that were singly tethered were stretched such that they overlapped during forward flow. Partial compaction began around 10 s after Dps arrived in the overlap region. Once forward flow ended, compaction was completed in around 4 seconds. The tip of the lower DNA molecule was then crosslinked to the base of the upper molecule and could not compact fully.

**Figure S3**

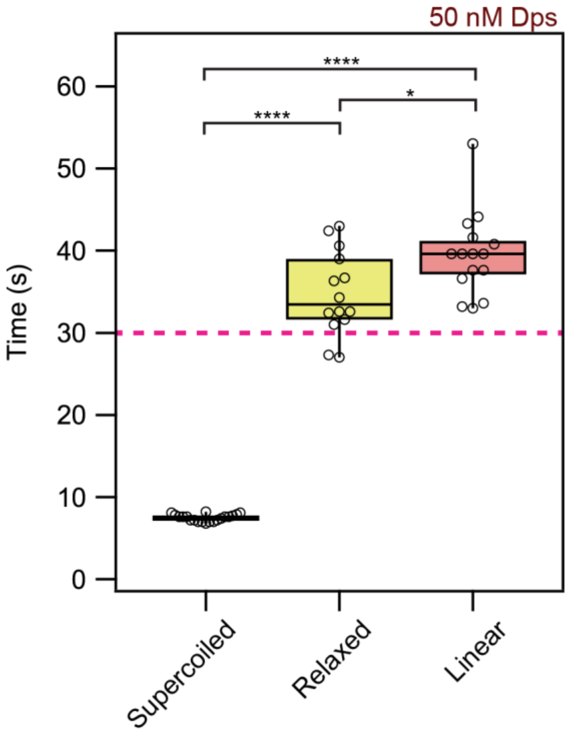


**Figure S3. Boxplots showing DNA compaction times at reduced SxO concentration.** We reduced the SxO concentration from 50 nM to 20 nM, decreasing the density of supercoiling in double tethered molecules without any nicks. Buffer flow containing 50 nM inlabeled Dps at 100 μL/minute was halted after 30 seconds (red dashed line). Boxplots showing the distribution of timepoints when 50 nM unlabeled Dps fully compacted the DNA in each conformation.

**Figure S4**

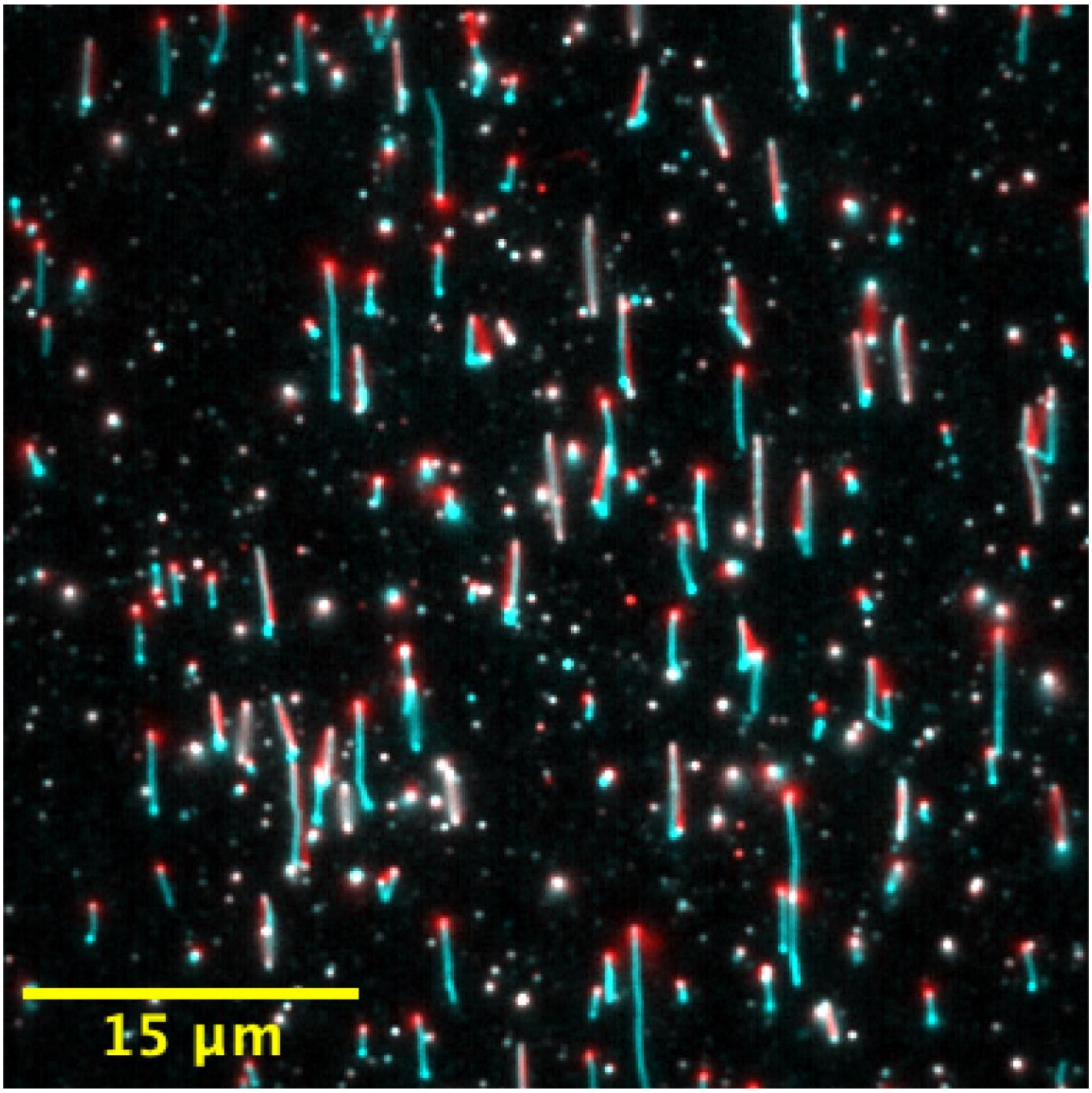


**Figure S4. PEG condensation of DNA under flow.** In order to observe condensation through a Dps-independent mechanism, we supplemented the imaging buffer with 3 mM $MgCl_2$ and additional PEG (8k), raising the PEG fractional volume to 20%. This high PEG buffer was introduced to the flow cell to induce compaction. Pictured is a 20-frame average of DNA molecules before addition of the high PEG buffer (red) and after the buffer exchange and the cessation of flow (cyan). The high PEG concentration in the buffer induced compaction in some molecules, but often bound singly tethered DNA to the surface in an extended conformation. No preference for the condensation of supercoiled DNA was observed.

**Figure S5**

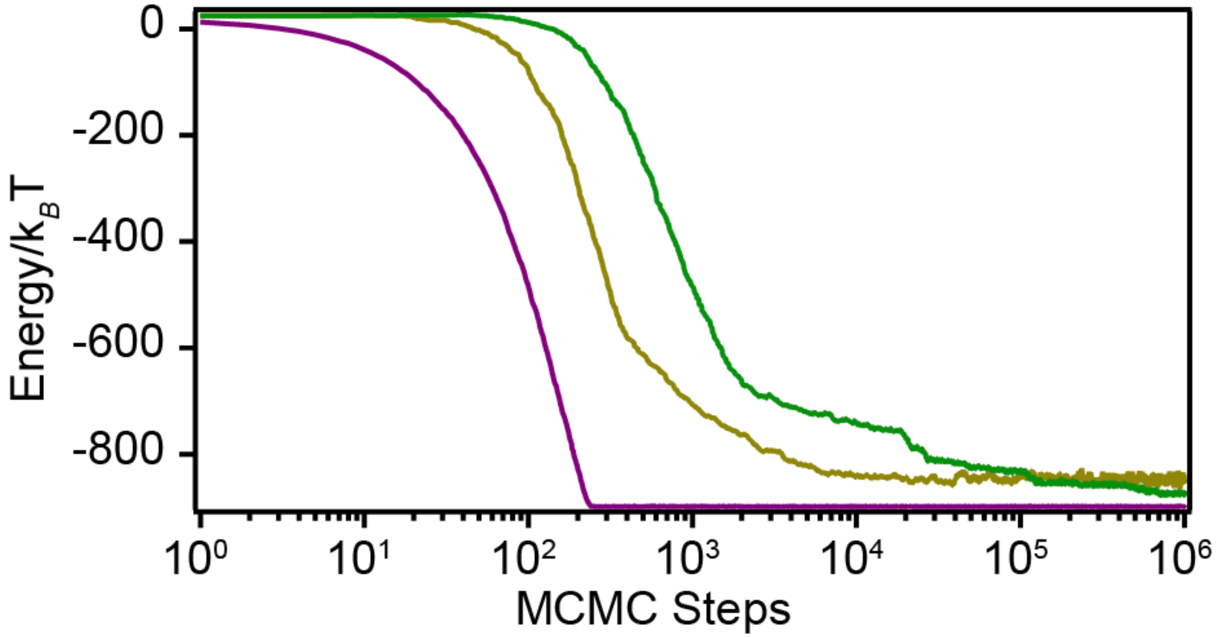


**Figure S5. MCMC simulations minimize free energy.** Each scenario in the MCMC simulations was run for $10^6$ steps and repeated five times. The total energy was averaged over each repeated run. In all cases the number of steps was sufficient to plateau at a minimum energy level. The first scenario (non-cooperative binding, violet curve) rapidly approached a minimum energy after <300 steps. The second scenario (nearest neighbor interactions only, yellow curve) approached a minimum energy after $<10^5$ steps. The third scenario (long range interactions, green curve) approached a minimum energy at the end of the simulation. To ensure that the true minimum had been reached, we also simulated the third scenario for $10^7$ steps and found the minimum to be stable.

**Figure S6**

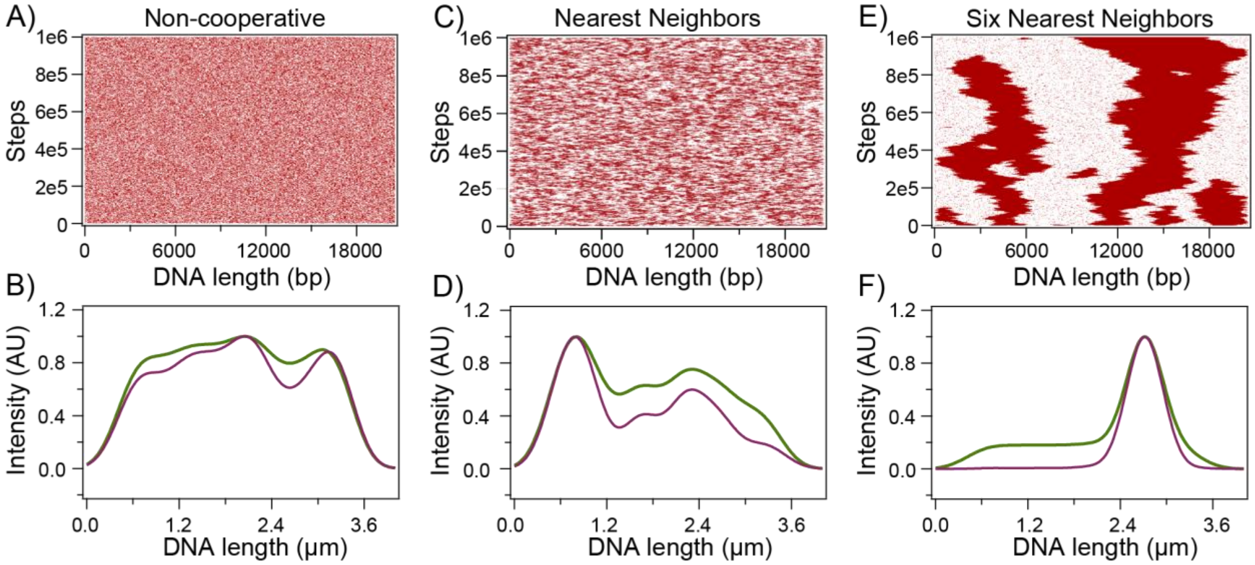


**Figure S6. Markov Chain Monte Carlo (MCMC) simulations of Dps-DNA complex formation at reduced binding energy. (A)** An MCMC simulation was performed where each Dps 12-mer was attracted to the DNA by a chemical potential of 3 $k_BT$ for binding DNA but with no cooperative interactions between neighboring 12-mers. Red dots represent DNA bound to Dps, and white patches represent unbound DNA. **(B)** Simulated fluorescence plot of a representative distribution of Dps (red) and DNA (green) seen in the simulation from **(A). (C)** A simulation as in **(A)** but with 0 $k_BT$ chemical potential for binding DNA and a 3 $k_BT$ interaction between any 12-mers bound in adjacent sites. **(D)** Simulated fluorescence plot of a representative distribution of Dps (red) and DNA (green) seen in the simulation from (B). **(E)** A simulation as in (A) but with 0 $k_BT$ chemical potential for binding DNA and a 0.5 $k_BT$ interaction between up to six adjacent 12-mers. **(F)** Simulated fluorescence plot of a representative distribution of Dps (red) and DNA (green) seen in the simulation from **(E)**.

**Figure S7**

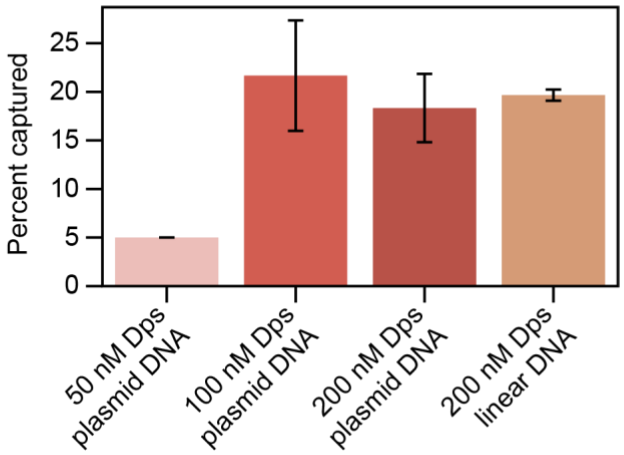


**Figure S7. Capture efficiency of secondary DNA by pre-formed Dps-DNA complexes.** Dps was introduced to flow cells at a range of concentrations (50-200 nM) and allowed to condense tethered DNA molecules. The Dps was then washed out of the flow cell with >10 volumes of imaging buffer. Additional DNA molecules (20.6 kb pSupercos lambda, either as an intact plasmid or a linearized plasmid) were then introduced into the flow cell and allowed to bind. The fraction of complexes that captured additional DNA molecules within a six-minute window was recorded and averaged over three separate experiments. The capture efficiency decreased for complexes formed at the lowest Dps concentrations.

**Supplementary Video SV1. Dps condensation of different topologies of DNA.** DNA strands labeled with SxO were immobilized on the coverslip. After a period of ~20 seconds, 200 nM Dps was flowed into the imaging chamber, causing the DNA to condense.

**Supplementary Video SV2. Dps condensation of different topologies of DNA.** Detailed view of a supercoiled (Y-shaped) DNA molecule condensed by Dps.

**Supplementary Video SV3. Dps condensation of different topologies of DNA.** Detailed view of a torsionally relaxed (J-shaped) DNA molecule condensed by Dps.
